# Development of thalamus mediates paternal age effect on offspring reading: A preliminary investigation

**DOI:** 10.1101/2020.05.20.105759

**Authors:** Zhichao Xia, Cheng Wang, Roeland Hancock, Maaike Vandermosten, Fumiko Hoeft

**Author notes:** **Corresponding Author:** Fumiko Hoeft, Department of Psychological Sciences and Brain Imaging Research Center, University of Connecticut, 850 Bolton Rd #1271, Storrs, CT 06269;.

## Abstract

The importance of (inherited) genetic impact in reading development is well-established. *De novo* mutation is another important contributor that is recently gathering interest as a major liability of neurodevelopmental disorders, but has been neglected in reading research to date. Paternal age at childbirth (PatAGE) is known as the most prominent risk factor for *de novo* mutation, which has been repeatedly shown by molecular genetic studies. As one of the first efforts, we performed a preliminary investigation of the relationship between PatAGE, offspring’s reading, and brain structure in a longitudinal neuroimaging study following 51 children from kindergarten through third grade. The results showed that greater PatAGE was associated significantly with worse reading, explaining an additional 9.5% of the variations after controlling for a number of confounds—including familial factors and cognitive-linguistic reading precursors. Moreover, this effect was mediated by volumetric maturation of the left posterior thalamus from ages 5 to 8. Complementary analyses indicated the PatAGE-related thalamic region was most likely located in the pulvinar nuclei and related to the dorsal attention network by using brain atlases, public datasets, and offspring’s diffusion imaging data. Altogether, these findings provide novel insights into neurocognitive mechanisms underlying the PatAGE effect on reading acquisition during its earliest phase and suggest promising areas of future research.

**Highlights:**

- Paternal age at childbirth (PatAGE) is negatively correlated with offspring’s reading abilities.
- PatAGE is related to volumetric maturation of the left posterior thalamus.
- Thalamic development mediates the PatAGE effect on reading.
- The PatAGE-related thalamic area is more likely to connect with the dorsal attention network.

## Introduction

There has been a global trend of postponed childbearing, especially in developed countries (Kohler, Billari, & Ortega, 2002). This so-called “postponement transition” is primarily owing to changing patterns of education, employment, and marriage (Khandwala, Zhang, Lu, & Eisenberg, 2017; Sobotka, 2010). Although the research field is still in its infancy, increasing evidence reveals that advanced paternal age at childbirth (PatAGE) increases risks for a wide range of neuropsychiatric conditions, such as schizophrenia and autism spectrum disorders (D’Onofrio et al., 2014).

In contrast to mental health, few studies investigated the effects of PatAGE on offspring’s cognition, such as reading, which is essential to success in modern society. The pioneering study in 1978 reported a negatively skewed distribution of PatAGE in 48 boys with reading disorders (RD; a.k.a. and referred here to dyslexia) (Jayasekara & Street, 1978). Four decades later, the topic remains underinvestigated and the existing findings are controversial. In a broader sample of 7-year-old children, Saha and colleagues revealed a significantly negative effect of PatAGE on offspring’s reading after controlling maternal age at childbirth (MatAGE), gestational age, sex, and race (Saha et al., 2009). However, when parental education and number of siblings were added to the statistical model, the effect of PatAGE on reading was no longer significant (Edwards & Roff, 2010). Such inconsistency underlies the importance of more research that controls for possible confounds examining the PatAGE effect on reading.

In addition to the controversy over the PatAGE effect on reading, no studies have yet examined the underlying mechanisms. Nascent research in molecular genetics, however, show that PatAGE explains nearly all the variance in the amount of *de novo* mutation, which is an alteration in a gene as the result of a mutation in a germ cell (egg or sperm) that increases by cell divisions of the gametes (approximately 38-fold in males at the age of 50 compared to females) (Breuss et al., 2019; Jónsson et al., 2017; Kong et al., 2012). Hence, *de novo* mutation is the most likely molecular mechanism underlying the PatAGE effect. In a separate line of research, *de novo* mutation is known to increase risk by up to 20-fold in neurodevelopmental disorders (De Rubeis et al., 2014; Deciphering Developmental Disorders Study, 2017). Taken together, it is conceivable that *de novo* mutations are at least partially responsible for the negative effect of PatAGE on offspring’s mental health, offering a plausible explanation of the PatAGE effect on children’s reading abilities.

At the neurocognitive level, whether and how cognitive-linguistic factors mediate the PatAGE effect on reading development is unknown. Studies to date have focused on genetic and environmental factors that contribute to the multifactorial liability of dyslexia (Pennington, 2006; Petrill, Deater-Deckard, Thompson, DeThorne, & Schatschneider, 2006). One such example, phonological processing, is thought to exert its effect more dominantly through inherited genetic impact, often estimated by family reading history (van Bergen, Bishop, van Zuijen, & de Jong, 2015). Under the same framework, whether PatAGE serves as a contributor to the multifactorial liability, and if so, what the neural and cognitive mediators (that would likely be heritable but not inherited traits if caused by *de novo* mutation) are, have not been examined. Brain measures derived from neuroimaging techniques, including magnetic resonance imaging (MRI), are particularly informative as they have been suggested as mediators between genetic etiology and behavioral outcome, acting as endophenotypes (Grasby et al., 2020; Shaw et al., 2012). Further, longitudinal investigation, combined with cross-sectional analysis, can provide comprehensive insights into the neural basis underlying typical reading acquisition and its impairment (Clark et al., 2014; Yeatman, Dougherty, Ben-Shachar, & Wandell, 2012).

Therefore, we conducted a preliminary study examining behavioral and multimodal neuroimaging data cross-sectionally and longitudinally in a cohort of 51 children followed from kindergarten through third grade in conjunction with analyses of publicly available datasets. The objective of the study was threefold: (1) to examine the relationship between PatAGE and offspring’s reading while systematically controlling for potential contributing/confounding factors; (2) to examine the role of previously known cognitive-linguistic precursors, neuroanatomy, and its maturational process in relation to PatAGE and reading; and (3) to understand the functional role of the neuroanatomical findings in this study by identifying convergent evidence through the use of brain atlases, public datasets, and offspring’s diffusion imaging data.

## Materials and Methods

### Participants

Participants in this study were drawn from a longitudinal NIH-funded project (K23HD054720) focusing on children’s reading development and followed from kindergarten (time-point 1 [*t*1], mean age = 5.58 years, *SD* = 0.43) to third grade (time-point 2 [*t*2], mean age = 8.30 years, *SD* = 0.46). All children were healthy native English speakers without neurological/psychiatric disorders (e.g., attention deficit/hyperactivity disorder) or contraindications to MRI based on parental reports. Among the participating children, 76% were White, 6% were Asian, and 18% were of multiradical heritage. About 8% of the children identified as Hispanic or Latino. Based on the annual household income, parental educational levels, and occupation, the participants in this study were of relatively high socioeconomic status (also see Black et al., 2012). The initial sample consisted of 51 children recruited from local newspapers, school mailings, flyers, and mothers’ clubs. In the behavior analyses, eight children were excluded because of attrition (n = 5), no record of PatAGE (n = 1), or being siblings (n = 2). In the latter case, the child with poorer T1 image quality was excluded. The final sample for behavioral analyses included 43 unrelated children (17 girls). In the neuroanatomical analysis, another 7 children (2 girls) were excluded because of incomplete T1 data collection or poor image quality at either time-point. In the diffusion imaging analysis, a sub-group of 23 children (8 girls) with the same acquisition sequence was included. The differences in either familial or behavioral measures between the entire and sub-groups were non-significant (all *p*’s > 0.1). The Institutional Review Boards of Stanford University where data were collected and the principal investigator was at the time of the study, and the University of California San Francisco where data were analyzed due to transition of the principal investigator, approved the present study. Both informed assent and consent were obtained from children and their guardians.

### Behavioral measurements

Demographics, family information, and performance on behavioral tests of the participants are summarized in Table 1. Family information collected at *t*1 includes: PatAGE; MatAGE; Adult Reading History Questionnaire from both parents (PatARHQ, MatARHQ) that were used to estimate familial history of reading difficulty (Lefly & Pennington, 2000); numbers of older and younger siblings; parental education (PatEDU, MatEDU); socioeconomic status (SES), a composite index computed from annual family income, parental education, and occupation with principal component analysis (Noble, Wolmetz, Ochs, Farah, & McCandliss, 2006); Home Observation Measurement of the Environment (HOME), an index for home environment including home literacy environment (Segers, Damhuis, de Sande, & Verhoeven, 2016). A battery of behavioral tests measuring intelligence, language, and reading-related skills was administrated. Verbal Comprehension, Concept Formation, and Visual Matching sub-tests of the Woodcock-Johnson III Tests of Cognitive Abilities (McGrew & Schrank, 2007) were used to estimate general cognitive abilities. These tests have reliabilities of at least 0.80 and have been used as a proxy for intelligence quotient (IQ) (Shaw, 2010). Vocabulary was measured with Peabody Picture Vocabulary Test (4^th^ edition) (Dunn & Dunn, 2007). Blending, Elision, Memory for Digit, Nonword Repetition sub-tests from the Comprehensive Test of Phonological Processing (CTOPP 1^st^ Edition) (Wagner, Torgesen, & Rashotte, 1999) were used to measure phonological skills. Rapid Automatized Naming (RAN; Objects and Colors sub-tests) (Wolf & Denkla, 2005) and Letter Identification sub-test of Woodcock Reading Mastery Test R/NU (WRMT-R/NU) (Mather, 1998) were also administered.

**Table 1.** Demographic profiles, familial variables, and performance on reading-related tests (n = 43). Acronyms: ARHQ, Adult Reading History Questionnaire; CS, composite score; CTOPP BW, Comprehensive Test of Phonological Processing, Blending sub-test; CTOPP EL, Comprehensive Test of Phonological Processing, Elision sub-test; CTOPP MD, Comprehensive Test of Phonological Processing, Memory for Digit sub-test; CTOPP NR, Comprehensive Test of Phonological Processing, Nonword Repetition sub-test; HOME, Home Observation Measurement of the Environment; Mat, maternal; Pat, paternal; PPVT, Peabody Picture Vocabulary Test; RAN COL, Rapid Naming, Colors sub-test; RAN LTR, Rapid Naming, Letters sub-test; RAN NUM, Rapid Naming, Numbers sub-test; RAN OBJ, Rapid Naming, Objects sub-test; RS, raw score; SES, socioeconomic status; SS, standard score; TOWRE PDE, Test of Word Reading, Phonemic Decoding Efficiency sub-test; TOWRE SWE, Test of Word Reading, Sight Word Efficiency sub-test; WJA RF, Woodcock-Johnson III Tests of Achievement, Reading Fluency sub-test; WJA SP, Woodcock-Johnson III Tests of Achievement, Spelling sub-test; WJC CF, Woodcock-Johnson III Tests of Cognitive Abilities, Concept Formation sub-test; WJC NR, Woodcock-Johnson III Tests of Cognitive Abilities, Numbers Reversed sub-test; WJC VC, Woodcock-Johnson III Tests of Cognitive Abilities, Verbal Comprehension sub-test; WJC VM, Woodcock-Johnson III Tests of Cognitive Abilities, Visual Matching sub-test; WRMT LID, Woodcock Reading Mastery Test, Letter Identification sub-test; WRMT PC, Woodcock Reading Mastery Test, Passage Comprehension sub-test; WRMT WA, Woodcock Reading Mastery Test, Word Attack sub-test; WRMT WID, Woodcock Reading Mastery Test, Word Identification sub-test.

|  | *Mean* | *Std.Dev* | *Min* | *Max* |
| --- | --- | --- | --- | --- |
| Time-point 1 |  |  |  |  |
| Age (years) | 5.58 | 0.43 | 5.03 | 6.99 |
| Gender, Male (%) | 60.50 | -- | -- | -- |
| Handedness, Right (%) | 88.40 | -- | -- | -- |
| WJC VC (SS) | 121.53 | 13.43 | 86 | 145 |
| WJC CF (SS) | 118.16 | 11.43 | 94 | 137 |
| WJC VM (SS) | 105.65 | 11.83 | 72 | 127 |
| WJC NR (SS) | 112.05 | 12.06 | 83 | 138 |
| PPVT (SS) | 121.23 | 9.94 | 97 | 148 |
| # Older Siblings (RS) | 0.65 | 0.81 | 0 | 3 |
| # Younger Siblings (RS) | 0.63 | 0.62 | 0 | 2 |
| PatAGE (years) | 36.12 | 4.91 | 24.78 | 46.71 |
| MatAGE (years) | 33.01 | 4.09 | 23.04 | 41.08 |
| PatARHQ (RS) | 0.35 | 0.14 | 0.09 | 0.66 |
| MatARHQ (RS) | 0.31 | 0.15 | 0.07 | 0.67 |
| PatEDU (years) | 16.95 | 2.05 | 13 | 22 |
| MatEDU (years) | 16.97 | 2.04 | 12 | 22 |
| SES (CS) <sup>a</sup> | 0.04 | 1.00 | -2.87 | 2.38 |
| HOME (RS) <sup>b</sup> | 51.39 | 2.25 | 44 | 55 |
| CTOPP BW (SS) <sup>c</sup> | 12.28 | 1.80 | 8 | 17 |
| CTOPP EL (SS) <sup>c</sup> | 11.93 | 2.80 | 7 | 19 |
| CTOPP MD (SS) <sup>c</sup> | 10.79 | 2.22 | 7 | 16 |
| CTOPP NR (SS) <sup>c</sup> | 11.23 | 2.85 | 6 | 19 |
| RAN OBJ (SS) | 100.56 | 17.57 | 55 | 135 |
| RAN COL (SS) | 97.98 | 16.51 | 55 | 137 |
| WRMT LID (SS) | 109.84 | 10.84 | 80 | 138 |
| Time-point 2 |  |  |  |  |
| Age (years) | 8.30 | 0.46 | 7.51 | 9.76 |
| WJC VC (SS) | 116.07 | 10.18 | 92 | 144 |
| WJC CF (SS) | 118.21 | 12.29 | 97 | 150 |
| WJC VM (SS) | 99.23 | 15.04 | 77 | 138 |
| WJC NR (SS) | 109.26 | 15.13 | 80 | 140 |
| PPVT (SS) | 120.02 | 14.56 | 81 | 160 |
| TOWRE SWE (SS) | 111.49 | 12.29 | 86 | 138 |
| TOWRE PDE (SS) | 106.35 | 14.85 | 77 | 144 |
| WRMT WID (SS) | 116.23 | 11.66 | 94 | 139 |
| WRMT WA (SS) | 113.84 | 14.40 | 90 | 146 |
| WRMT PC (SS) | 114.91 | 9.95 | 99 | 141 |
| WJA RF (SS) | 112.49 | 16.31 | 84 | 162 |
| WJA SP (SS) | 105.28 | 18.33 | 74 | 148 |
| CTOPP BW (SS) <sup>c</sup> | 12.67 | 2.24 | 6 | 16 |
| CTOPP EL (SS) <sup>c</sup> | 13.00 | 3.11 | 4 | 17 |
| CTOPP MD (SS) <sup>c</sup> | 10.47 | 2.60 | 5 | 15 |
| CTOPP NR (SS) <sup>c</sup> | 10.09 | 2.26 | 6 | 16 |
| RAN NUM (SS) | 100.35 | 12.47 | 76 | 129 |
| RAN LTR (SS) | 102.98 | 11.63 | 78 | 134 |
| RAN OBJ (SS) | 96.53 | 17.07 | 62 | 132 |
| RAN COL (SS) | 97.02 | 15.29 | 60 | 121 |

Notes: <sup>a</sup> SES: $n = 38$ ; <sup>b</sup> HOME: $n = 41$ ; <sup>c</sup> T Scores are presented for CTOPP sub-tests where mean is 10 and SD is 3. All other test scores are in standard scores where the mean is 100 and SD is 15.

The same set of tests were used at *t*2 (Table 1). Numbers and Letters sub-tests of RAN were further included to measure print-sound mapping efficiency. Additionally, we administered tests measuring different aspects of reading, including Sight Word Efficiency and Phonemic Decoding Efficiency sub-tests from the Test of Word Reading Efficiency (TOWRE 1^st^ Edition) (Torgesen, Wagner, & Rashotte, 1999), Word Identification, Word Attack, and Passage Comprehension sub-tests from WRMT-R/NU, and Reading Fluency and Spelling sub-tests from WJ-III Tests of Achievement.

### Image acquisition

High-resolution T1-weighted images were collected at both time-points with the following parameters: 128 slices; thickness = 1.2 mm; NEX = 1; repetition time = 8.5 ms; echo time = 3.4 ms; inversion time = 400 ms; in-plane resolution = 256 × 256; voxel size = 0.9 × 0.9 × 1.2 mm^3^; flip angle = 15 °; field of view = 22 cm. High-angular resolution diffusion-imaging data were collected at *t*2 with the following parameters: 46 axial slices; slice thickness = 3 mm; repetition time = 5000 ms; echo time = 81.7 ms; in-plane resolution = 128 × 128; voxel size = 2.0 × 2.0 × 3.0 mm^3^; 150 directions with b = 2500 s/mm^2^; 6 volumes with b = 0 s/mm^2^. All images were acquired using a GE Healthcare 3.0 T 750 scanner with eight-channel phased-array head coil at Richard M. Lucas Center for Imaging at Stanford University. The quality of images was qualitatively evaluated by an investigator who was blinded to the behavioral and demographic information prior to any analyses.

### Behavior analyses

To reduce the dimensionality of behavioral metrics, factor analyses were conducted on reading-related tests for each time-point; *t*1: Blending, Elision, Memory for Digits, Nonword Repetition sub-tests of CTOPP, Objects and Colors sub-tests of RAN, Letter Identification sub-test of WRMT; *t*2: Blending, Elision, Memory for Digits, Nonword Repetition sub-tests of CTOPP, Numbers, Letters, Objects and Colors sub-tests of RAN, Sight Word Efficiency and Phonemic Decoding Efficiency sub-tests of TOWRE, Word Identification, Word Attack, Passage Comprehension sub-tests of WRMT-R/NU, Reading Fluency and Spelling sub-tests of WJ-III Tests of Achievement. The Maximum Likelihood, Varimax, and Bartlett methods were used for extraction, rotation, and factor score calculation. The criterion of eigenvalue greater than 1 was used to identify factors. From *t*1 behavioral metrics, we obtained two factors, explaining 53.8% of the total variance. Since phonological awareness (PA) and RAN loaded heavily on each factor, we named these two factors *t*1PA and *t*1RAN (Fig. 1A). Given that PA and RAN are the most reliable predictors for reading development in alphabetic languages (Caravolas et al., 2012), we used these two composite scores as cognitive-linguistic precursors of reading in subsequent analyses. Using the same approach, we extracted three factors from *t*2 behavioral metrics, explaining 67.2% of the total variance. The scores were labeled as *t*2READ, *t*2PA, and *t*2RAN according to the corresponding factor loading (Fig. 1B).

**Fig. 1.**
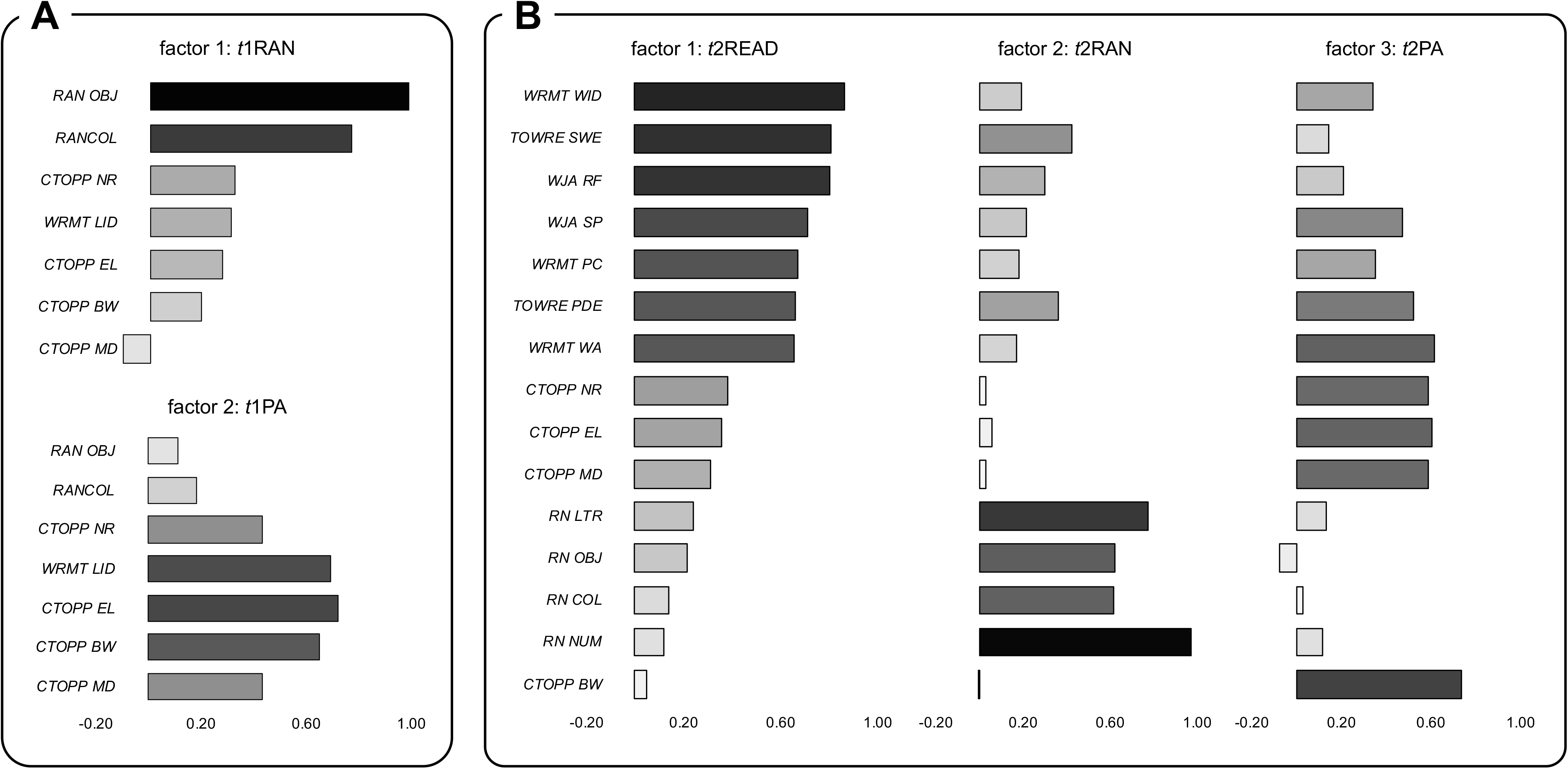
Principal components that extracted from reading-related tests. **A**. Component loadings for each factor at time-point 1. **B**. Component loadings for each factor at time-point 2. *Acronyms: CTOPP BW, Comprehensive Test of Phonological Processing, Blending sub-test; CTOPP EL, Comprehensive Test of Phonological Processing, Elision sub-test; CTOPP MD, Comprehensive Test of Phonological Processing, Memory for Digit sub-test; CTOPP NR, Comprehensive Test of Phonological Processing, Nonword Repetition sub-test; RAN COL, Rapid Naming, Colors sub-test; RAN LTR, Rapid Naming, Letters sub-test; RAN NUM, Rapid Naming, Numbers sub-test; RAN OBJ, Rapid Naming, Objects sub-test; t1, time-point 1; t2, time-point 2; TOWRE PDE, Test of Word Reading, Phonemic Decoding Efficiency sub-test; TOWRE SWE, Test of Word Reading, Sight Word Efficiency sub-test; WJA RF, Woodcock-Johnson III Tests of Achievement, Reading Fluency sub-test; WJA SP, Woodcock-Johnson III Tests of Achievement, Spelling sub-test; WRMT LID, Woodcock Reading Mastery Test, Letter Identification sub-test; WRMT PC, Woodcock Reading Mastery Test, Passage Comprehension sub-test; WRMT WA, Woodcock Reading Mastery Test, Word Attack sub-test; WRMT WID, Woodcock Reading Mastery Test, Word Identification sub-test*.

Since a consensus on the definition of advanced paternal age remained lacking (Couture, Delisle, Mercier, & Pennings, 2020), in this study we treated PatAGE as a continuous variable rather than separating children into different PatAGE groups. To examine the relationship between PatAGE and reading, we first calculated the zero-order correlation between them. Once the correlation was significant, hierarchical linear regressions were conducted to answer four questions in the following order: (1) whether the PatAGE effect on reading remains significant after controlling for demographic variables and general intelligence; (2) whether the PatAGE effect on reading exists after additionally regressing out MatAGE, which is known to correlate highly with PatAGE and is a possible confound; (3) whether the PatAGE effect on reading is present above and beyond familial risk (representing inherited risk; Swagerman et al., 2017), and environmental factors to address the issue of multifactorial liability; (4) whether the PatAGE effect on reading is explained by *t*1 cognitive-linguistic precursors of reading to examine whether the most common predictors were the mediating factor.

Specifically, in the first model we entered *t*2 age, sex, handedness, and average performance IQ (pIQ) across *t*1 and *t*2 in the first step and PatAGE in the second step (Table 2). In the second model, besides the aforementioned nuisance variables, we regressed out MatAGE, which was found correlated with both PatAGE and *t*2READ. In the third model, familial risk measured by ARHQ of both parents (van Bergen et al., 2015), and environmental factors including educational level of both parents (Edwards & Roff, 2010), number of older and younger siblings (Price, 2008), SES (Pan et al., 2016), HOME (Segers et al., 2016), which are known to be associated with reading were additionally controlled. In the final model, *t*1PA and *t*1RAN were entered in the fourth step, just before PatAGE, to examine whether the PatAGE effect was present beyond *t*1 cognitive-linguistic skills. Given that *t*1RAN and *t*1PA did not correlate with PatAGE (both *r*’s < 0.01; Table S1), these two cognitive-linguistic precursors were not examined further for the mediating relationship. All statistics were done with SPSS 24.0 (IBM, Inc.), and *p-*values were two-tailed while statistical significance was set at 0.05.

**Table 2.**
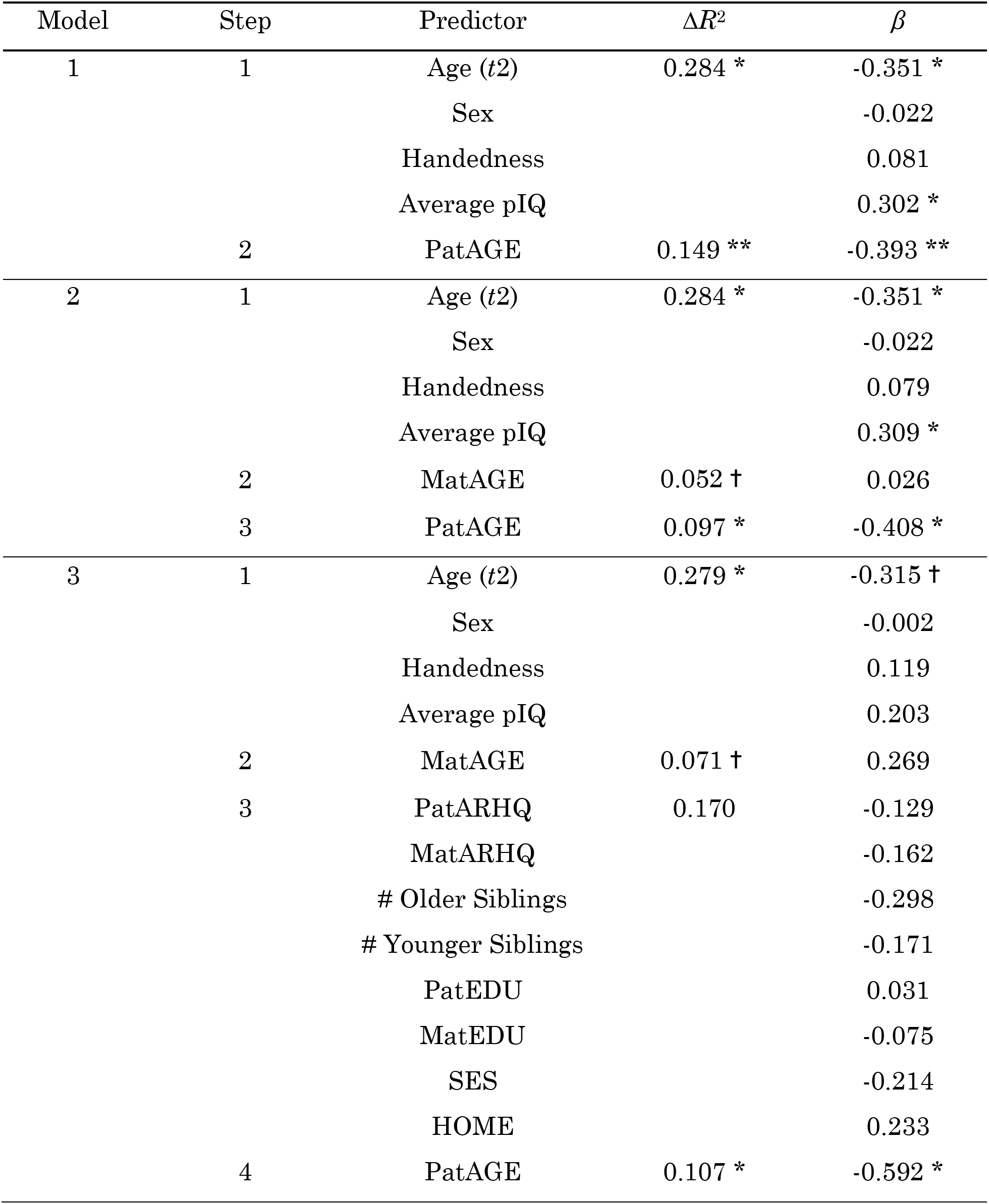
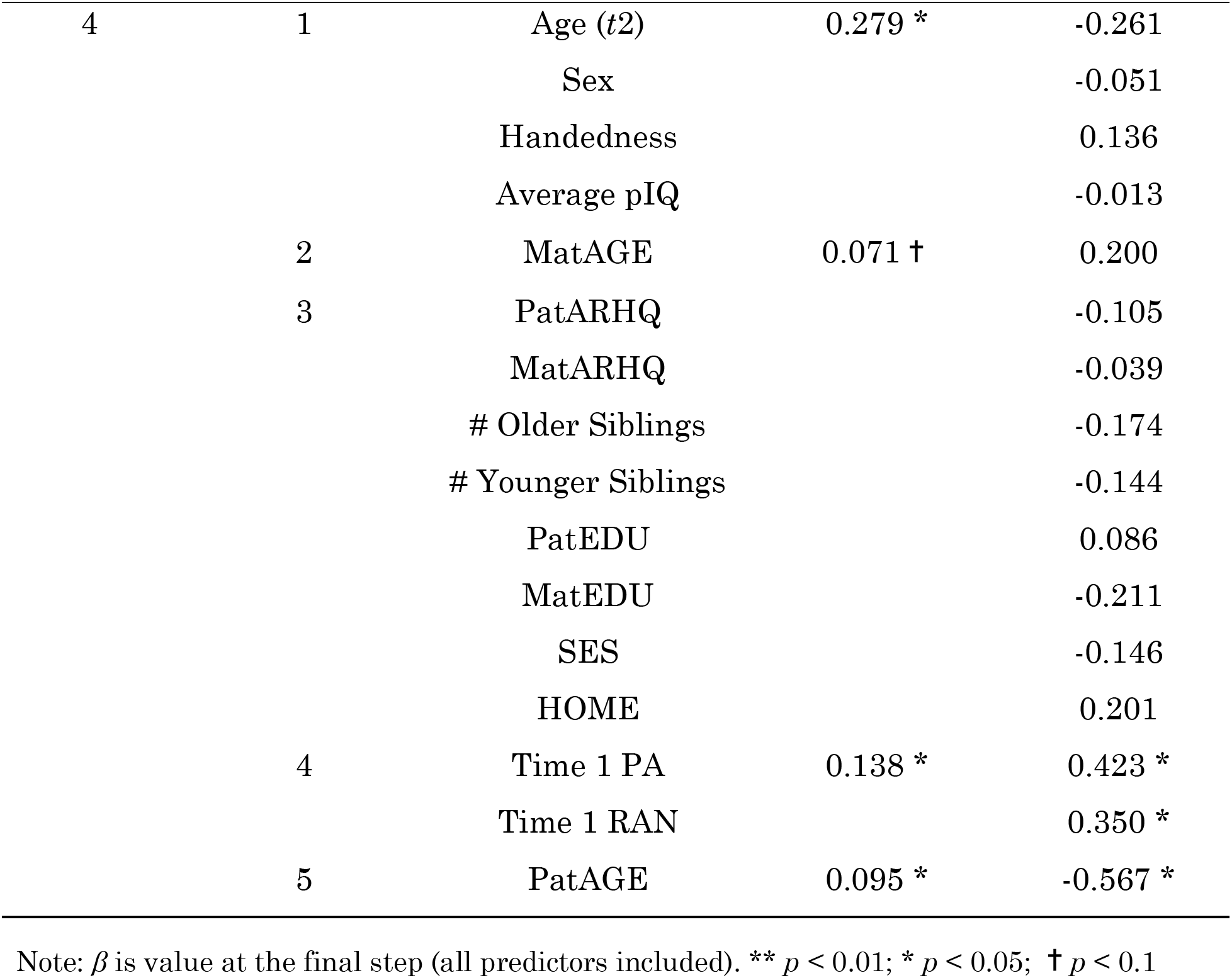
Results of multiple linear regression analyses examining the unique contribution of paternal age on offspring’s reading at time-point 2. *Acronyms: ARHQ, Adult Reading History Questionnaire; EDU, educational level; HOME, Home Observation Measurement of the Environment; Mat, maternal; PA, phonological awareness; Pat, paternal; pIQ, performance intelligence quotient; RAN, rapid naming; SES, socioeconomic status; t1, time-point 1; t2, time-point 2*.

| Model | Step | Predictor | $\Delta R^2$ | $\beta$ |
| --- | --- | --- | --- | --- |
| 1 | 1 | Age (*t2*) | 0.284 * | -0.351 * |
|  |  | Sex |  | -0.022 |
|  |  | Handedness |  | 0.081 |
|  |  | Average pIQ |  | 0.302 * |
|  | 2 | PatAGE | 0.149 ** | -0.393 ** |
| 2 | 1 | Age (*t2*) | 0.284 * | -0.351 * |
|  |  | Sex |  | -0.022 |
|  |  | Handedness |  | 0.079 |
|  |  | Average pIQ |  | 0.309 * |
|  | 2 | MatAGE | 0.052 † | 0.026 |
|  | 3 | PatAGE | 0.097 * | -0.408 * |
| 3 | 1 | Age (*t2*) | 0.279 * | -0.315 † |
|  |  | Sex |  | -0.002 |
|  |  | Handedness |  | 0.119 |
|  |  | Average pIQ |  | 0.203 |
|  | 2 | MatAGE | 0.071 † | 0.269 |
|  | 3 | PatARHQ | 0.170 | -0.129 |
|  |  | MatARHQ |  | -0.162 |
|  |  | # Older Siblings |  | -0.298 |
|  |  | # Younger Siblings |  | -0.171 |
|  |  | PatEDU |  | 0.031 |
|  |  | MatEDU |  | -0.075 |
|  |  | SES |  | -0.214 |
|  |  | HOME |  | 0.233 |
|  | 4 | PatAGE | 0.107 * | -0.592 * |
| 4 | 1 | Age (*t*2) | 0.279 * | -0.261 |
|  |  | Sex |  | -0.051 |
|  |  | Handedness |  | 0.136 |
|  |  | Average pIQ |  | -0.013 |
|  | 2 | MatAGE | 0.071 † | 0.200 |
|  | 3 | PatARHQ |  | -0.105 |
|  |  | MatARHQ |  | -0.039 |
|  |  | # Older Siblings |  | -0.174 |
|  |  | # Younger Siblings |  | -0.144 |
|  |  | PatEDU |  | 0.086 |
|  |  | MatEDU |  | -0.211 |
|  |  | SES |  | -0.146 |
|  |  | HOME |  | 0.201 |
|  | 4 | Time 1 PA | 0.138 * | 0.423 * |
|  |  | Time 1 RAN |  | 0.350 * |
|  | 5 | PatAGE | 0.095 * | -0.567 * |

Note: $\beta$ is value at the final step (all predictors included). \*\* $p < 0.01$ ; \* $p < 0.05$ ; † $p < 0.1$

### Structural image preprocessing

Both cross-sectional and longitudinal analyses were conducted by using the voxel-based morphometry toolbox (v435; http://www.neuro.uni-jena.de/vbm/) with SPM8 (v4667; http://www.fil.ion.ucl.ac.uk/spm8/) implemented in Matlab R2014a. In the cross-sectional data preprocessing for *t*1 and *t*2, individual T1 volumes were segmented into gray matter, white matter, and cerebrospinal fluid with a resampling at 1.5 mm^3^. Then, the gray matter segments were registered to a T1 template in Montreal Neurological Institute (MNI) space by using both affine normalization and Diffeomorphic Anatomical Registration Through Exponentiated Lie Algebra (Ashburner, 2007), and subsequently modulated by the “affine and non-linear” modulation (http://www.neuro.uni-jena.de/vbm/segmentation/modulation/). The modulated images containing regional tissue volume of gray matter for each voxel were smoothed with an 8-mm full-width half-maximum isotropic Gaussian kernel. Voxels with gray matter values < 0.1 were excluded to avoid edge effects.

In the longitudinal data preprocessing, the “Preprocessing of Longitudinal Data” module that contains specific preprocessing steps for processing longitudinal structural MRI data was used. Intra-subject realignment, bias correction, segmentation, and normalization were conducted sequentially as described elsewhere (Ridgway et al., 2007). After applying spatial smoothing with an 8-mm full-width half-maximum Gaussian kernel, we obtained maps of gray matter volume for both time-points. We then generated a brain map reflecting gray matter volume (GMV) change from *t*1 to *t*2 for each child (such that a positive value indicates enlarging from *t*1 to *t*2).

### Whole-brain analyses

Prior to voxel-wise analyses, we examined relationships between PatAGE and global measurement (i.e., the total intracranial volume; defined as the sum of total gray matter, white matter, and cerebrospinal fluid) at each time-point (*t*1TIV and *t*2TIV). Then, we examined whether PatAGE correlated with the change of TIV from *t*1 to *t*2 (ΔTIV) while controlling for the baseline measure (*t*1TIV). To explore relationships between regional GMV at each time-point (i.e., cross-sectional analyses), as well as the change of regional GMV (ΔGMV) across time-points (*t*2GMV-*t*1GMV) with PatAGE (i.e., longitudinal analysis), voxel-wise whole-brain regressions were conducted while controlling for global measurements. Specifically, *t*1TIV or *t*2TIV was controlled in cross-sectional analyses for *t*1 and *t*2, respectively. In the longitudinal analysis, *t*1TIV and ΔTIV were controlled to exclude the effects from initial gross volume and its development. Since correlations between *t*1TIV, ΔTIV, and PatAGE were not significant (all *p*’s > 0.1), the models were free from multicollinearity. Topological Family Wise Error (FWE) correction was used to determine the corrected thresholds of statistical significance. All clusters significant at a threshold of *p*-cluster < 0.05 corrected for the whole brain (*p*-voxel < 0.001 for height) were reported in MNI space. Region-of-interest (ROI) analyses were conducted in the significant clusters to examine the robustness and specificity of the effect. For this, values of each voxel in the cluster were extracted and averaged, then included in hierarchical multiple regression analyses as the dependent variable. Demographic variables (*t*1 or *t*2 age, sex, handedness, average pIQ across *t*1 and *t*2 for cross-sectional data; *t*1 age, time interval between *t*1 and *t*2, sex, handedness, average pIQ across *t*1 and *t*2 for longitudinal data) and global measurements (*t*1TIV or *t*2TIV for cross-sectional data; *t*1TIV and ΔTIV for longitudinal data) were entered in the first step, while PatAGE was entered in the second step. Then, we further controlled for MatAGE and MatARHQ since they showed significant correlations with PatAGE as in previous analyses of this study.

### Mediation analyses

One of the main objectives of this study was to investigate possible neurocognitive mediators of the PatAGE effect on reading. At the brain level, two analytical strategies were used. For the primary approach, we conducted whole-brain analyses on *t*2READ cross-sectionally and longitudinally in the same way as we did for PatAGE. Next, we administered conjunction analysis to identify overlapping areas that showed significant associations with both PatAGE and *t*2READ, following which the mediation relationship was examined. Alternatively, we took an ROI approach if no significant cluster survived multiple correction at the whole-brain level on *t*2READ. Specifically, we examined the relationship between ΔGMV and children’s *t*2READ in the cluster significantly associated with PatAGE (hereafter PatAGE-cluster). The partial correlation coefficient was calculated while controlling for demographic variables (*t*2 age, sex, handedness, average pIQ across *t*1 and *t*2), global measurements (*t*1TIV and ΔTIV), and cognitive-linguistic precursors (*t*1PA and *t*1RAN) that were significantly associated with *t*2READ in previous regression analysis of this study. Once a region was identified in either whole-brain or ROI analysis, we examined whether the PatAGE effect on reading was mediated by brain measures. The model was adjusted for demographic variables (*t*2 age, sex, handedness, average pIQ across *t*1 and *t*2), global measurements (*t*1TIV, ΔTIV), and *t*1 cognitive-linguistic precursors (*t*1RAN and *t*1PA). Bootstrapping (10,000 samples) was used to obtain 95% confidence interval of the indirect effect. If the confidence interval does not contain zero, a significant indirect effect is indicated.

Existing evidence suggests that dyslexia is largely genetically transmitted from parents (often assayed by parent-report of reading difficulty) (Soden et al., 2015; Swagerman et al., 2017). Further, twin studies find a dissociation between sources of variance in phonological and orthographic processes, with variance in phonological skills being primarily genetic compared to orthographic skills (Olson, Wise, Conners, Rack, & Fulker, 1989; Olson et al., 2011). These findings are consistent with the idea that PA mediates the negative effect of parental reading difficulty (a proxy for inherited genetic transmission) on reading in offspring (van Bergen et al., 2015). In line with the previous literature, we observed significant correlations between MatARHQ and *t*1PA, MatARHQ and *t*2READ, *t*1PA and *t*2READ. We therefore examined the role of PA on the relationship between the history of maternal reading difficulty and lower reading performance in offspring. Demographic variables (age at *t*2, sex, handedness, average pIQ across *t*1 and *t*2) and the other cognitive-linguistic precursor (*t*1RAN) were controlled statistically. If both a PatAGE effect (a proxy for non-inherited genetic risk) and MatARHQ effect via PA processing (a proxy for inherited genetic risk) are to be observed in the current samples, this study then supports the multifactorial liability model.

PROCESS procedure (release 2.16.1) implemented in SPSS was used to conduct these mediation analyses (Hayes, 2013).

### Complementary analyses

We adopted multiple complementary analytical approaches to depict fine-grained spatial localization and connectivity patterns of the PatAGE-cluster, capitalizing on the fact that these have been shown to inform possible functional roles of a particular brain region (in this case, the left posterior thalamus) in the absence of a comprehensive set of cognitive measures. First, we spatially localized the PatAGE-cluster with two brain atlases. (1) Given that the thalamus consists of multiple nuclei with different functions, we calculated the percentage of overlapped voxels between the PatAGE-cluster and each thalamic nucleus from the Morel Atlas, a histological atlas that is optimal for thalamic targets in MNI space (Jakab, Blanc, Berényi, & Székely, 2012; Krauth et al., 2010); (2) Given that the connectivity pattern provides information about the function of a given brain region (Barron, Eickhoff, Clos, & Fox, 2015), we used Oxford Thalamic Connectivity Probability Atlas (https://fsl.fmrib.ox.ac.uk/fsl/fslwiki/Atlases) with the atlasquery tool implemented in FSL to obtain the probability that the PatAGE-cluster is structurally connected to different cortical areas.

To further identify the possible functional roles of the PatAGE-cluster and complement the results from analyses using the histological and connectivity probability atlases, we examined PatAGE-cluster-associated cortical patterns by using online databases provided in Neurosynth v0.5 (Yarkoni, Poldrack, Nichols, Van Essen, & Wager, 2011). In particular, we generated a co-activation map by including all fMRI studies in the database (N > 10,900), with the PatAGE-cluster as ROI. False Discovery Rate (FDR) corrected *q* < 0.01 was used as the threshold to obtain significant regions reported in fMRI studies when the PatAGE-cluster is also reported (i.e., forward inference). In addition, we generated a map of whole-brain resting-state functional connectivity (RSFC) by using the 1000 Functional Connectome dataset (Biswal et al., 2010). The center of gravity of the PatAGE-cluster (MNI: x = −19.6, y = −28.1, z = 6.9) was used as the seed, and its connectivity to the rest of the brain was calculated. The resultant brain map was thresholded with a liberal cutoff value of *r* = 0.01, the same as in the previous literature (Yang, Rosenblau, Keifer, & Pelphrey, 2015). To be conservative, we only considered the overlapping regions between the co-activation map and the RSFC map as the cortical pattern of the PatAGE-cluster. Sørensen-Dice coefficients between the conjunction map and the 7 large-scale intrinsic connectivity networks (i.e., visual, somatomotor, dorsal attention, ventral attention, limbic, frontoparietal, and default networks) from Yeo et al. (2011) were calculated to examine which functional network most overlapped with the PatAGE-cluster-associated cortical pattern. Here we used an adult network template because studies in Neurosynth that were used to produce the co-activation map were conducted in participants with a wide range of ages and the 1000 Functional Connectome dataset mainly consists of adult data.

Given that the functional community structure in children is to some extent different from that in adults and the highest uncertainty was found in attention networks (Tooley, Bassett, & Mackey, 2021; Vijayakumar et al., 2021), in the final step, we examined the structural connectivity pattern in a sub-group of our own samples where diffusion imaging data were available. Moreover, we adopted a pediatric intrinsic functional network template that was extracted from 670 children aged 9-11 years (Tooley et al., 2021) with the same approach as in Yeo et al. (2011). Specifically, we analyzed white matter connectivity, where fibers passing the PatAGE-cluster were reconstructed using deterministic tractography. Diffusion-weighted imaging preprocessing was performed by using ExploreDTI (Leemans, Jeurissen, Sijbers, & Jones, 2009). Next, to visually inspect for possible artefacts, rigorous motion correction with CATNAP and eddy current correction were conducted by using the required reorientation of the b-matrix (Leemans & Jones, 2009). The diffusion tensors were calculated using a non-linear regression procedure (Pierpaoli & Basser, 1996). The individual datasets were non-rigidly normalized to MNI space. Next, whole-brain tractography was performed for each individual dataset using a deterministic approach (Basser, Pajevic, Pierpaoli, Duda, & Aldroubi, 2000). Fibers (streamlines) were reconstructed by defining seed points distributed uniformly throughout the data at 2.0 × 2.0 × 2.0 mm^3^ resolution, following the main direction with the step size set at 1.0 mm. Fiber tracking was discontinued when the fiber entered a voxel with fractional anisotropy < 0.2 or made a high angular turn (angle > 40°) or when the fiber was outside the fiber length range of 50-500 mm. Two analyses were then conducted: (1) To localize fibers and get a general view, the PatAGE-cluster was used as ROI and all fibers passing through the ROI were delineated. The delineated fibers and their projection points were visually inspected, after which individual maps were binarized and summed to acquire the probabilistic map across participants. (2) To complement the Neurosynth analysis above and identify the functional network most relevant to the PatAGE-cluster, the number of streamlines passing through the PatAGE-cluster and each of the 7 template pediatric functional networks (Tooley et al., 2021) was calculated and normalized by dividing global density of the target network (percentage of total voxels). The results were treated as the connectivity strength and compared between candidate networks achieved in the previous analyses. Furthermore, we examined the correlations between the connectivity strength with PatAGE and *t*2READ.

## Results

### PatAGE is negatively associated with offspring’s reading above and beyond commonly known predictors

PatAGE (*M* = 36.12 years, *SD* = 4.91, *Range* = 25-47; Table 1; Fig. S1A) was positively correlated with MatAGE (*r* = 0.63, *p* = 5 × 10^-6^) but not with other potentially confounding demographic variables reported in the past such as SES, number of siblings and parental education (all *p’*s > 0.1; Table S1). As to the main objective of this study, greater PatAGE was significantly correlated with lower reading composite scores in offspring (*t*2READ; *r* = −0.39, *p* = 0.011). Similar to PatAGE and not surprisingly, MatAGE was negatively correlated with *t*2READ (*r* = −0.33, *p* = 0.031). No significant correlations were found between PatAGE and cognitive-linguistic skills typically found to be predictors of later reading ability at either time-point (*p*’s > 0.1, for *t*1PA, *t*1RAN, *t*2PA, and *t*2RAN). In accordance with prior literature on factors that predict reading outcomes (Segers et al., 2016; Thompson et al., 2015; van Bergen et al., 2015), lower *t*2READ was predicted by poorer reading reported by mothers (MatARHQ; *r* = −0.46, *p* = 0.002), poorer home literacy environment measured by HOME (*r* = 0.31, *p* = 0.047), and poorer cognitive-linguistic skills at time-point 1 (*t*1PA: *r* = 0.46, *p* = 0.002; *t*1RAN: *r* = 0.31; *p* = 0.041).

To examine whether the PatAGE effect on reading existed above and beyond commonly identified confounds and additional variables known to influence reading development, hierarchical linear regressions were conducted with *t*2READ as the dependent variable in a systematic and hypothesis-driven fashion. In the first model, before PatAGE was entered, demographic variables (*t*2 age, sex, handedness) and general intelligence (average pIQ across two time-points) were entered as predictors in the first step. The negative effect of advanced PatAGE remained significant, explaining an additional 14.9% of the variance (*t* = −3.12, *p* = 0.004; Model 1 in Table 2). In the second model, MatAGE was further added in the second step, since it was significantly correlated with PatAGE and *t*2READ. As shown in Table 2 (Model 2), PatAGE explained an additional 9.7% of the variance in reading outcomes (*t* = −2.48, *p* = 0.018). Then, in the third model, familial risk (PatARHQ, MatARHQ) and environmental factors (number of siblings, parental education, SES, and HOME) in relation to reading development were added. We still observed a significant PatAGE effect, explaining an additional 10.7% of the variance (*t* = − 2.45, *p* = 0.023; Model 3 in Table 2). Thus far, we demonstrated that the PatAGE effect on reading was not accounted for by factors that predict children’s reading outcomes and known to be either inherited or environmental. In the final model, we investigated its relationship with early cognitive-linguistic predictors of reading outcomes by entering *t*1PA and *t*1RAN in the fourth step. The PatAGE effect on offspring’s reading was above and beyond that of cognitive-linguistic variables, explaining an additional 9.5% of the variance (*t* = −2.71, *p* = 0.014; Model 4 in Table 2; Fig. S1B). In accord with the prior literature, *t*1PA and *t*1RAN also significantly predicted *t*2READ in the final model and accounted for 13.8% of the variation (*t*1PA: *t* = 2.87, *p* = 0.010; *t*1RAN: *t* = 2.19, *p* = 0.042). That is, contributions from PatAGE and cognitive-linguistic precursors were relatively independent, and they jointly predicted children’s reading outcomes.

### PatAGE is associated with thalamic maturation

In the brain analyses, we did not observe significant correlations between PatAGE and TIV at *t*1 (*r* = −0.27, *p* = 0.109), *t*2 (*r* = −0.29, *p* = 0.085), or ΔTIV from *t*1 to *t*2 (*r* = −0.11, *p* = 0.537). Second, whole-brain analyses on regional GMV at each time-point did not show any significant clusters at the FWE corrected threshold of *p*-cluster < 0.05 (*p*-voxel < 0.001 for height). Finally, we examined the PatAGE effect on regional ΔGMV while controlling for *t*1TIV and ΔTIV. Results revealed a significantly positive correlation between PatAGE and ΔGMV in a cluster covering the left posterior thalamus (i.e., PatAGE-cluster; *p* = 0.015, FWE corrected, 367 voxels, 1,239 mm^3^, peak MNI coordinate [-27, −30, 6]; Fig. 2A). Specifically, greater PatAGE was associated with less GMV decrease. To verify that this effect was not due to confounds, hierarchical linear regression analyses were performed. In the first model, after regressing out nuisance variables commonly controlled in longitudinal VBM analysis (*t*1 age, time interval between *t*1 and *t*2, sex, handedness, average pIQ across *t*1 and *t*2, *t*1TIVand ΔTIV), PatAGE explained 35.2% of the ΔGMV variance of the PatAGE-cluster (*t* = 4.71, *p* < 0.001). Since MatARHQ and MatAGE were significantly correlated with PatAGE, we additionally included them as covariates in the second model, and found PatAGE still explained 17.1% of the ΔGMV variance of the PatAGE-cluster (*t* = 3.18, *p* = 0.004; Fig. 2B).

**Fig. 2.**
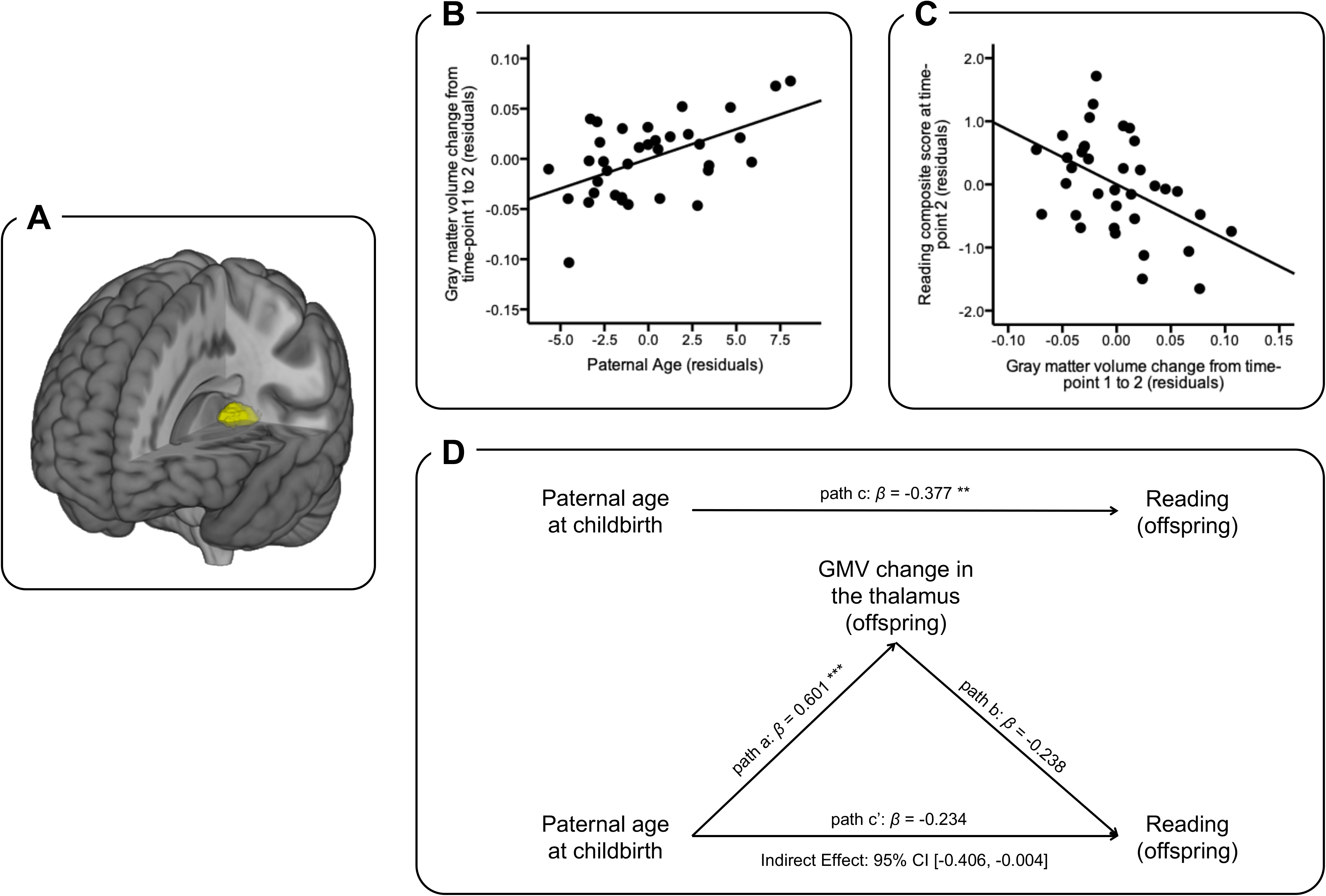
Results of the whole-brain longitudinal voxel-based morphometry and region-of-interest (ROI) analysis. **A.** Brain region significantly correlated with paternal age (the yellow cluster; defined as ROI). **B**. Scatter plot of the relationship between gray matter volume change in the ROI and paternal age. The linear regression line is presented. **C**. Scatter plot of the relationship between gray matter volume change in the ROI and composite score of reading at time-point 2. The linear regression line is presented. **D.** The effect of paternal age on offspring’s reading is mediated by gray matter volume change in the thalamus. Confounds were controlled statistically. The bias-corrected 95% confidence interval for indirect effect was [-0.406, −0.004], indicating a significant mediation relationship. *** *p* < 0.001; ** *p* < 0.01

### The PatAGE effect on offspring’s reading is mediated by **Δ**GMV in the left posterior thalamus

First, no significant correlations were observed between *t*2READ with TIV at each time-point or ΔTIV from *t*1 to *t*2 (all *p*’s > 0.1). In either the whole-brain cross-sectional or longitudinal analyses, we did not find clusters showing GMV-reading correlations survived the FWE correction. Therefore, the subsequent analyses were conducted based on the PatAGE-cluster revealed in the previous step. After we found ΔGMV in the PatAGE-cluster was correlated with *t*2READ while nuisance variables (*t*2 age, sex, handedness, average pIQ across *t*1 and *t*2), global measurements (*t*1TIV, ΔTIV), and cognitive-linguistic precursors of reading (*t*1PA, *t*1RAN) were statistically controlled (partial *r* = −0.48, *p* = 0.011; Fig. 2C), we further examined the mediation relationship. As shown in Fig. 2D, ΔGMV significantly mediated the negative effect of advanced PatAGE on offspring’s reading; 95% confidence interval was [-0.406, −0.004] when nuisance variables (age at *t*2, sex, handedness, average pIQ across *t*1 and *t*2), global measurements (TIV at *t*1, ΔTIV), and cognitive-linguistic precursors (*t*1PA, *t*1RAN) were statistically controlled. These results are in contrast to the commonly found results in the literature that we also find in the present study, i.e., *t*1PA mediates the negative effect of family history on offspring’s reading (95% confidence interval was [-0.249, −0.001] when nuisance variables (age at *t*2, sex, handedness, average pIQ across *t*1 and *t*2) and the other cognitive-linguistic precursor (*t*1RAN) were controlled; Fig. S2).

### PatAGE-cluster is localized in the pulvinar nuclei and linked to the dorsal attention network

To understand the neurostructural profile of the PatAGE-cluster in the left thalamus, we compared it with a histological atlas and a connectivity atlas. 279 out of 367 voxels in the cluster overlapped with the human thalamus of the Morel histological atlas (Jakab et al., 2012; Krauth et al., 2010), while the remaining 88 voxels were unlabeled, possibly because the cluster also contained white matter. As presented in Fig. 3A, within the overlapping region, 215 voxels (76.9%) were in the subdivision labeled as pulvinar nuclei, especially the medial portion, known to have widespread connections with the inferior parietal lobule (Arcaro, Pinsk, & Kastner, 2015). These results were corroborated by examining the Thalamic Connectivity Probability Atlas (http://fsl.fmrib.ox.ac.uk/fsl/fslview/atlas.html): the PatAGE-cluster was most likely localized in the subdivision that was connected to the posterior parietal cortex, with a probability of 48.2% (Fig. 3B).

**Fig. 3.**
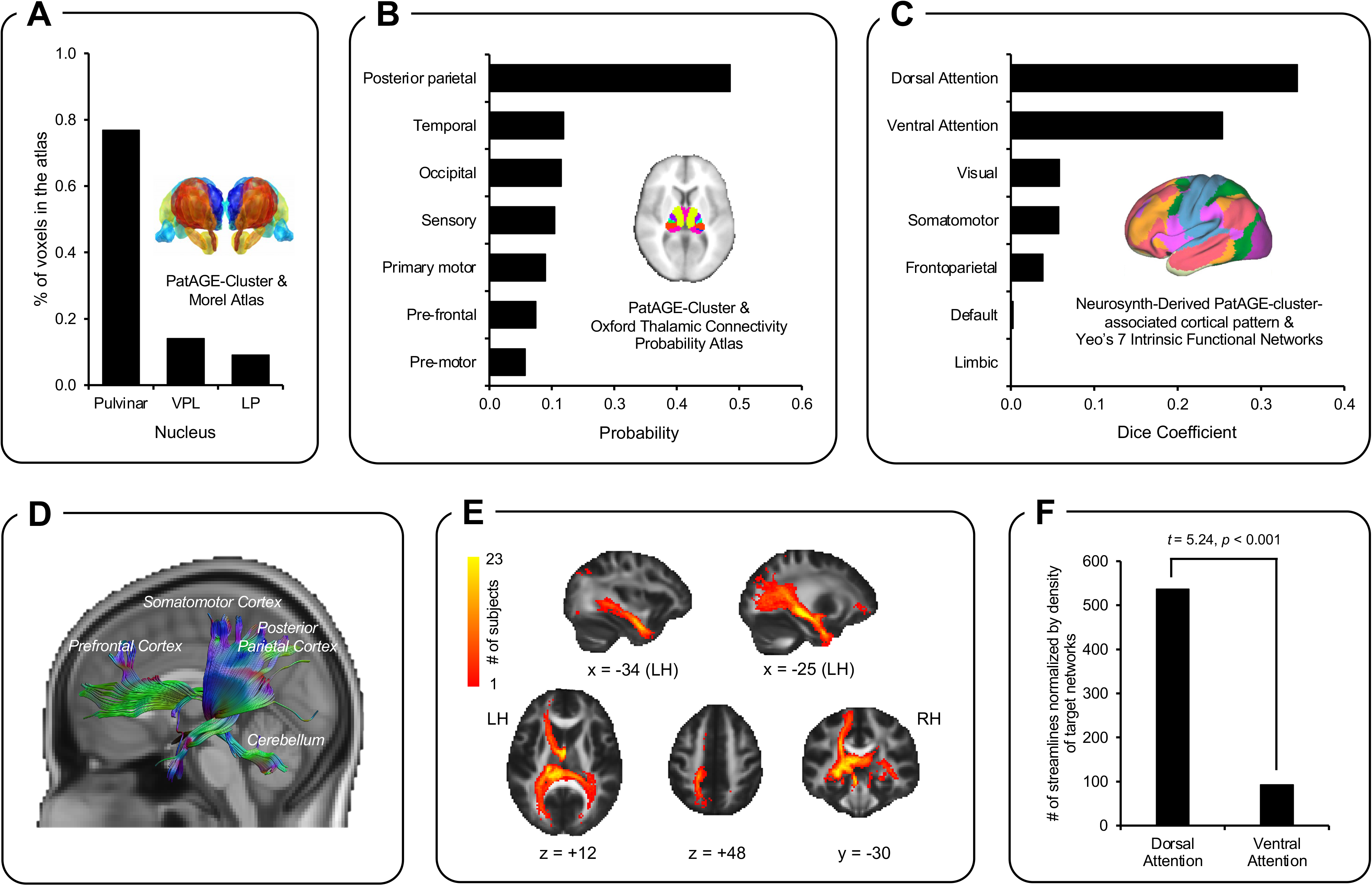
Results of the complementary analyses on the PatAGE-cluster (i.e., the left posterior thalamus) with atlases, public database, and white matter tractography of the PatAGE-related thalamic region using subject-specific diffusion imaging data. **A.** Bar plot displaying the percentage of total voxels in the PatAGE-cluster overlaps with divisions of the Morel Atlas (https://www.lead-dbs.org/helpsupport/knowledge-base/atlasesresources/atlases/). **B.** Bar plot showing the probability of the cluster belonging to different subdivisions of the Oxford Thalamic Connectivity Probability Atlas (https://fsl.fmrib.ox.ac.uk/fsl/fslwiki/Atlases), calculated by the “autoaq” function implemented in FSL. **C.** Bar plot showing the degree of overlap between the overlapping areas of Neurosynth-derived co-activation/resting-state functional connectivity maps and Yeo’s 7 intrinsic functional networks represented by Dice coefficients. Dice coefficient measures the similarity between the overlapping areas and a given function network, ranging from 0 to 1. While 0 indicates the two networks are disjoint, 1 indicates the two networks are identical. D. Example of reconstructed fibers in a representative child with the seed being the PatAGE-cluster. E. Intersection across 23 children with diffusion imaging data is shown for demonstrative purposes. The color bar represents the number of subjects where the streamline is observed in a given voxel. F. The DAN compared to the VAN derived from Tooley’s 7 intrinsic functional networks derived from pediatric data showed a significantly greater number of streamlines (normalized by global density of the target network [percentage of total voxels]) to go through the PatAGE-cluster. *Acronyms: CL, central lateral nucleus; CM, central median nucleus; LP, lateral posterior nucleus; VLpv, ventral lateral posterior nucleus, ventral; VPI, ventral posterior inferior nucleus; VPL, ventral posterior lateral nucleus; VPM, ventral posterior medial nucleus; LH, left hemisphere; RH, right hemisphere*.

Next, we examined the cortical pattern of the PatAGE-cluster by utilizing two approaches available in Neurosynth v0.5 (Yarkoni et al., 2011). These included the generation of a meta-analytic map of regions that co-activate with the PatAGE-cluster across more than 10,900 fMRI studies and an RSFC map from the PatAGE-cluster to the rest of the brain by using the 1000 Functional Connectome dataset (Biswal et al., 2010). The co-activated areas included subcortical structures and cortical regions such as bilateral intraparietal sulci, inferior temporal gyrus, and frontal eye fields in the frontal cortex (Fig. S3A). The RSFC map showed a similar but more widespread pattern than the co-activation map (Fig. S3B). A conjunction analysis revealed that the bilateral frontal eye fields, intraparietal sulci, middle temporal visual area (V5/MT), and cerebellum were among the overlapped regions across the two approaches, in addition to subcortical structures (Fig. S3C).

Sørensen-Dice coefficients (*s*) between the overlapping areas and the previously identified networks deriving from resting-state functional MRI data (Yeo et al., 2011) were calculated. The derived pattern of overlapping areas showed the greatest resemblance to the dorsal attention network (DAN; *s* = 0.344; Fig. 3C) and to the ventral attention network (VAN; *s* = 0.254), much higher than its resemblance to other networks (visual network: *s* = 0.058; somatomotor network: *s* = 0.057; limbic network: *s* < 0.001; frontoparietal network: *s* = 0.039; and default network: *s* = 0.003). Together with the aforementioned findings utilizing structural atlases, these results using large-scale fMRI databases from functional neuroimaging studies point to the attention network, particularly the DAN, to be the candidate brain functional system associated with the PatAGE-cluster in the left thalamus.

Finally, we analyzed diffusion imaging data available in a sub-group of 23 participants to confirm DAN was more likely the candidate system associated with the PatAGE effect on reading than VAN. Because the diffusion imaging data were collected at time-point 2, different from the previous analyses with Neurosynth, here we adopted Tooley’s functional network template that was extracted from data of children aged 9-11 years (Tooley et al., 2021). Using deterministic tractography, we reconstructed white matter fibers through the PatAGE-cluster, covering inferior fronto-occipital fasciculus, corticospinal tract, forceps major, superior corona radiata, as well as anterior and posterior limbs of the internal capsule. Fig. 3D shows reconstructed fibers in a representative child and Fig. 3E shows intersection across participants, for demonstrative purposes. In line with previous findings of this study, the PatAGE-cluster showed significantly stronger connectivity (defined as the total number of streamlines divided by global density of the target network [percentage of total voxels]) with DAN than with VAN (*t* = 5.24, *p* < 0.001; Fig. 3F). No significant correlations were found between PatAGE-cluster-Network streamlines and PatAGE or *t*2READ (all *p*’*s* > 0.1).

## Discussion

In this study, we observed a significantly negative effect of advanced PatAGE on offspring’s reading at the earliest stages of formal schooling from ages 5 to 8, independent of confounds (e.g., maternal age) and factors that play key roles in learning to read (i.e., family reading history, environmental factors, and cognitive-linguistic precursors of reading), explaining an additional 9.5% of the variance. Furthermore, we revealed volumetric maturation of the left thalamus as a potential neural endophenotype mediating this effect. We identified that this area is most relevant to the dorsal attention network with brain atlases, public datasets, and offspring’s diffusion imaging data. These findings contrast and complement the current literature linking phonological and orthographic processing in reading to the left temporo-parietal and occipito-temporal regions. The mediation revealed here was distinct from the mediating role of phonological processing on the relationship between reading and familial risk, which has been attributed to hereditary effects. Taken together, this study provides novel and converging evidence suggesting PatAGE as a significant factor that is associated with offspring’s reading, independent of phonological processing, possibly through the maturational process of the left posterior thalamus.

### The PatAGE effect on offspring’s reading

Jayasekara and Street (1978), for the first time, reported that advanced PatAGE was associated with a greater incidence of dyslexia, independent of SES and birth order. While the analysis was conducted in dyslexic boys, Saha and colleagues extended the finding to a broader population of 7-year-old children with varying reading abilities measured using Wide Range Achievement Test (Saha et al., 2009). Negative PatAGE effects on several cognitive measures, including reading, were observed after controlling for confounds that included MatAGE, SES, and parental psychiatric illness. A follow-up study re-analyzed the same dataset and found that the PatAGE effect was no longer significant after further adjusting parental education and the number of siblings (Edwards & Roff, 2010). Therefore, the PatAGE effect on reading was equivocal, and the inconsistency was related to covariates controlled in the model, especially parental characteristics such as educational level.

In the present study, with the range of PatAGE restricted to 25-47 years, we found PatAGE was negatively associated with reading performance measured using a variety of tests, even after additionally controlling for strong predictors of reading that were not included in previous studies (Edwards & Roff, 2010; Saha et al., 2009). These predictors included familial reading history and cognitive-linguistic skills (e.g., phonological processing) that shown to be more genetically than environmentally mediated, as well as home literacy environment (Hulme, Snowling, Caravolas, & Carroll, 2005; Swagerman et al., 2017; van Bergen et al., 2015). These findings support an adverse PatAGE effect on reading and suggest such effect may occur through a mechanism different to factors such as inherited genetic and environmental risks.

While the number of studies examining the PatAGE effect on reading is too few to infer the potential mechanisms, studies on PatAGE-linked neuropsychiatry disorders offer insights. One predominant explanation is that advanced PatAGE exerts its effect on the risk of disorders such as autism and schizophrenia through accumulated *de novo* genetic mutations and epigenetic modifications (e.g., DNA methylation and repressive histone modification) in paternal gametes (Deciphering Developmental Disorders Study, 2017; Girard et al., 2016; Saha et al., 2009).

At a more macroscopic scale, understanding of the mechanisms can be deepened by identifying intermediate (endo)phenotypes through behavioral and neuroimaging measures such as we did in the current study. That is, advanced PatAGE may impact precursors of neurodevelopmental disorders, which in turn leads to a higher occurrence of such disorders (Cannon, 2009). For example, the likelihood of having impaired social functioning in offspring, a core symptom of psychiatric disorders, increases with PatAGE (Weiser et al., 2008). While the underlying mechanisms are yet to be fully understood, multifactorial liability confers risk for neurodevelopmental disorders and may involve liability such as *de novo* mutations in addition to inherited and environmental risks. Adding to prior research, the current study offers insights into potential mechanisms at the macroscopic level.

### The intermediary role of the left posterior thalamus

The thalamus is an important relay center in the human brain, receiving information from sensory cortices and relaying it to higher-level association cortices. Previous studies paint a mixed picture on thalamic development: gross volume relative to its brain size is smaller (Sussman, Leung, Chakravarty, Lerch, & Taylor, 2016) or larger (Brain Development Cooperative, 2012) in older compared to younger children of ages 4 to 18, and the pulvinar compared to other thalamic nuclei show no apparent change with age (Raznahan et al., 2014). Despite controversial evidence on typical thalamic maturation, its anomalous development undoubtedly affects the growth of other cortical and subcortical brain regions (Ball et al., 2012), which in turn could impact higher-level cognitive processes that underlie typical reading acquisition. In support of this, anomalies in thalamic structure (Giraldo-Chica, Hegarty, & Schneider, 2015), activation (Diaz, Hintz, Kiebel, & von Kriegstein, 2012; Koyama, Molfese, Milham, Mencl, & Pugh, 2020), and connectivity (Müller-Axt, Anwander, & von Kriegstein, 2017; Paz-Alonso et al., 2018; Tschentscher, Ruisinger, Blank, Diaz, & von Kriegstein, 2018) are associated with dyslexia. While most of these studies adopt a cross-sectional design with adult participants, here we conducted a longitudinal investigation and found the volumetric change in the posterior thalamus from ages 5 to 8 was significantly associated with PatAGE; children with younger fathers showed GMV decrease, whereas those with older fathers showed less decrease or showed an increase. This pattern suggests that PatAGE is associated with the development of this subcortical structure. Noteworthy, although no significant PatAGE effect was observed when examining a single time-point, it does not indicate this effect cannot manifest at a specific age. Instead, it suggests that compared with brain morphometry at a particular time-point, PatAGE affects the maturation process more and underscores the importance of considering developmental dynamics when examining brain-behavior relationships. A similar pattern has been revealed in other neural measurements, such as white matter development in dyslexia (Yeatman et al., 2012).

Moreover, examination of the cluster’s location with the Morel Atlas suggested the foci in the left pulvinar, an integral region supporting visuo-spatial attention (Amso & Scerif, 2015; Fischer & Whitney, 2012), and attentional control (Barron et al., 2015; Xuan et al., 2016). Analysis with connectivity-based thalamic atlas showed that this region was most likely to overlap with the subdivision connected with posterior parietal areas. Furthermore, RSFC and co-activation maps produced by Neurosynth revealed connectivity patterns were suggestive of the attention networks, especially the DAN. Finally, analysis of children’s diffusion imaging-based connectivity with the pediatric functional network template was also suggestive of the DAN. Studies have repeatedly demonstrated the relationship between visuo-spatial attention and reading (Facoetti, Franceschini, & Gori, 2019; Vidyasagar & Pammer, 2010). First, visuo-spatial attention has been associated with acquiring orthographic knowledge (Stevens & Bavelier, 2012) and decoding skills (Matthews & Martin, 2015). Further, both dyslexic adults and children show deficits in visuo-spatial attention, such as impaired motion perception, lower visuo-spatial span capacities, slower responses during visuo-spatial attention-orienting tasks, and local precedence on global perception (Bosse, Tainturier, & Valdois, 2007; Buchholz & Davies, 2008; Franceschini, Bertoni, Gianesini, Gori, & Facoetti, 2017; Gori, Seitz, Ronconi, Franceschini, & Facoetti, 2015). Longitudinal research also demonstrate impaired visuo-spatial processing in pre-reading kindergarteners as a causal risk factor of future poor reading (Bertoni, Franceschini, Ronconi, Gori, & Facoetti, 2019; Carroll, Solity, & Shapiro, 2015; Franceschini, Bertoni, et al., 2017; Franceschini, Gori, Ruffino, Pedrolli, & Facoetti, 2012; S. Gori & Facoetti, 2015; Gori et al., 2015). Finally, targeted interventions such as action video game training and motion discrimination training effectively improve reading and reading-related cognitive skills in affected children via enhancing visuo-spatial attention and visual-to-auditory attentional shifting (Bertoni et al., 2019; Franceschini & Bertoni, 2019; Franceschini, Bertoni, et al., 2017; Franceschini et al., 2013; Franceschini, Trevisan, et al., 2017; Gori et al., 2015; Lawton, 2016). Together, these findings indicate that maturation of the pulvinar and brain networks underlying visuo-spatial attention are parsimonious neurocognitive mechanisms that may be impacted by advanced PatAGE, impeding reading acquisition. It should be noted that with the current data, we could not exclude the possibility it was the PatAGE-related individual difference in reading that influenced the maturation of thalamus and its connection with the attentional networks (Skeide et al., 2017). In fact, recent studies have revealed a bidirectional relationship between reading acquisition and the development of the underlying brain networks (Wang, Joanisse, & Booth, 2020; Wang, Pines, Joanisse, & Booth, 2021). Here we proposed that paternal age may influence thalamic development, which in turn affects reading acquisition. On the other hand, as children develop reading skills, the left posterior thalamus may in turn be impacted. Further studies are needed to examine this hypothesis.

To date, research investigating the PatAGE effect on brain networks and corresponding cognitive processes is scarce. As the first step, Shaw et al. (2012) revealed PatAGE effects on cortical morphometry. However, the authors did not examine the relationship with cognitive functions, making the study somewhat inconclusive as to the role of PatAGE on neurocognitive processes. Taking one step further, the current study revealed thalamic maturation as an intermediary between PatAGE and reading—a specific behavioral phenotype, offering insights into the complex mechanisms underlying PatAGE effects.

### Limitations and future directions

In the present study, we observed a negative PatAGE effect on offspring’s reading and the left posterior thalamus as a possible brain intermediary. Given the preliminary nature of this investigation and the small sample size, the findings should be interpreted with caution until they are replicated in large independent samples. Second, PatAGE in this study was restricted to 25-47 years, with which we observed a negative linear relationship. The findings hence do not necessarily extend to children with extremely young and old fathers. For example, Saha et al. (2009) observed a non-linear relationship with a range of PatAGE between 14 to 66 years, while the relationship appears to be linear in the age range as ours. Of relevance, young fatherhood is also associated with adverse outcomes in offspring but possibly due to other factors, including immature sperm and economic disadvantages (Chen et al., 2008). Third, the complementary analyses implied DAN as the candidate functional system associated with PatAGE. It should be noted that the brain atlases and public datasets implemented in Neurosynth are primarily from research on adults, while children have specific characteristics regarding brain organization (Tooley et al., 2021; Vijayakumar et al., 2021). We tempted to address this issue by adopting the functional network template for children in analyzing diffusion imaging data available in a subgroup of the current samples and found that the PatAGE-cluster more strongly connected with DAN than VAN. Nevertheless, the neurostructural profile and function of the PatAGE-cluster need to be re-visited as more pediatric-specific atlases and tools are available. Fourth, while we found that the left posterior thalamus mediated the PatAGE effect on reading, it remains unknown why this subcortical structure is susceptible to advanced PatAGE (and related to *de novo* mutations). Given that typical thalamic maturation is also affected by prenatal and postnatal factors such as preterm birth (Ball et al., 2012), questions including how PatAGE influences the growth of thalamus and relevant functional systems, together with other factors, require further elaboration.

To advance understanding of the PatAGE effect on reading, future studies are warranted in which a more comprehensive battery of behavioral tests (e.g., measuring visuo-spatial attention, executive function, etc.), neural measures (e.g., task-driven activation), and molecular approaches measuring the number and origins of *de novo* mutations (e.g., trio-based whole-genome/exome sequencing; Jin et al., 2017) are included. Fusing neural, cognitive, and molecular genetic approaches at multiple levels will provide the much-needed vertical and multi-level explanatory models that will further our understanding of risk factors associated with poor reading. In particular, future research aiming at disentangling different sources of genetic variations related to reading development and their interplays will greatly further our understanding. In addition, advanced research designs such as the intergenerational neuroimaging approach can be adopted to gain in-depth knowledge on how multiple factors, including PatAGE, affect the development of offspring’s reading and the corresponding networks interactively from preliteracy to mature stages of reading (Ho, Sanders, Gotlib, & Hoeft, 2016; Hoeft & Hancock, 2017).

## Conclusion

The current study examined the PatAGE effect on offspring’s reading at both behavioral and neurobiological levels. The results provide initial evidence that advanced PatAGE is a relatively independent factor associated with poor reading outcomes in beginning readers, above and beyond previously identified familial and cognitive-linguistic precursors. This effect was mediated by the maturation of the posterior thalamus, suggesting a neurobiological pathway to intergenerational influence on reading acquisition, complementing prior findings that offspring’s reading is influenced by parental reading via offspring’s phonological skills (van Bergen et al., 2015). Based on these findings, we argue that PatAGE should be regarded as an important factor influencing literacy development, and included in a cumulative risk (and protection) model (Hayiou-Thomas, Smith-Woolley, & Dale, 2020; Menghini et al., 2010; Pennington, 2006; van Bergen, van der Leij, & de Jong, 2014).

## Authors’ Contributions

F. Hoeft designed the study and collected data with her students. F. Hoeft, Z.C. Xia and C. Wang conceived the particular idea of the manuscript. Z.C. Xia, C. Wang and M. Vandermosten analyzed the data. Z.C. Xia, F. Hoeft, C. Wang, R. Hancock, and M. Vandermosten wrote and revised the manuscript.

## Acknowledgments

The authors thank all the families for their participation in this longitudinal study. They also thank Albert Galaburda and Tuong-Vi Nguyen for their thoughtful suggestions during manuscript preparation.

## Data Availability Statement

Data that support the findings of this study are available from the corresponding author on request.

## Abbreviations

ARHQ: Adult Reading History Questionnaire
CI: confidence interval
DAN: dorsal attention network
DNA: deoxyribonucleic acid
FDR: false discovery rate
FWE: family wise error
MatAGE: maternal age at childbirth
MNI: Montreal Neurological Institute
MRI: magnetic resonance imaging
PA: phonological awareness
PatAGE: paternal age at childbirth
pIQ: performance intelligence quotient
RAN: rapid automatized naming
RD: reading disorder
READ: reading composite score
ROI: region-of-interest
RSFC: resting-state functional connectivity
SES: socioeconomic status
t1: time-point 1
t2: time-point 2
TIV: total intracranial volume
V5/MT: middle temporal visual area
VAN: ventral attention network
VBM: voxel-based morphometry

