## Supplemental Materials for "Development of thalamus mediates paternal age effect on offspring reading: A preliminary investigation"

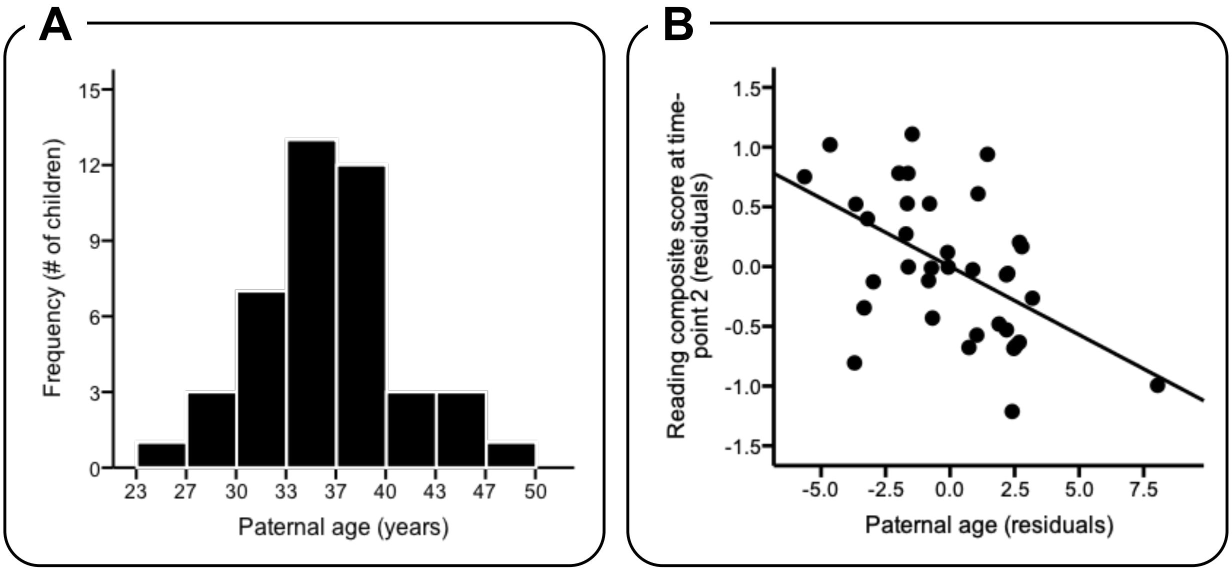


**Figure S1.** Distribution of paternal age at childbirth and its correlation with reading. **A.** Histogram of paternal age. **B.** Partial regression plot representing the correlation between paternal age and offspring's reading. Reading composite scores were calculated using factor analysis on reading-related tests at time-point 2 and adjusted for demographic variables. The linear regression line is presented.

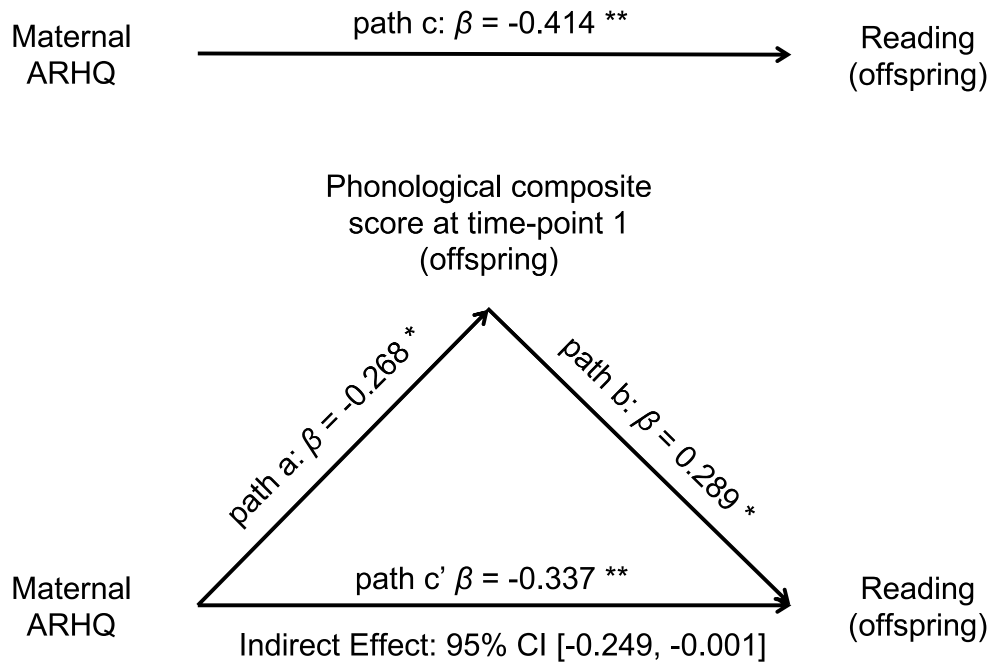


**Figure S2.** The effect of maternal reading history (measured by ARHQ) on offspring's reading at time-point 2 is mediated by the phonological composite score at time-point 1. Confounds were controlled statistically. The bias-corrected 95% confidence interval for indirect effect was [-0.249, -0.001], indicating a significant mediating relationship between familial history and offspring's reading. *Acronyms: ARHQ, Adult Reading History Questionnaire; CI, confidence interval;* ** $p < 0.01$; * $p < 0.05$

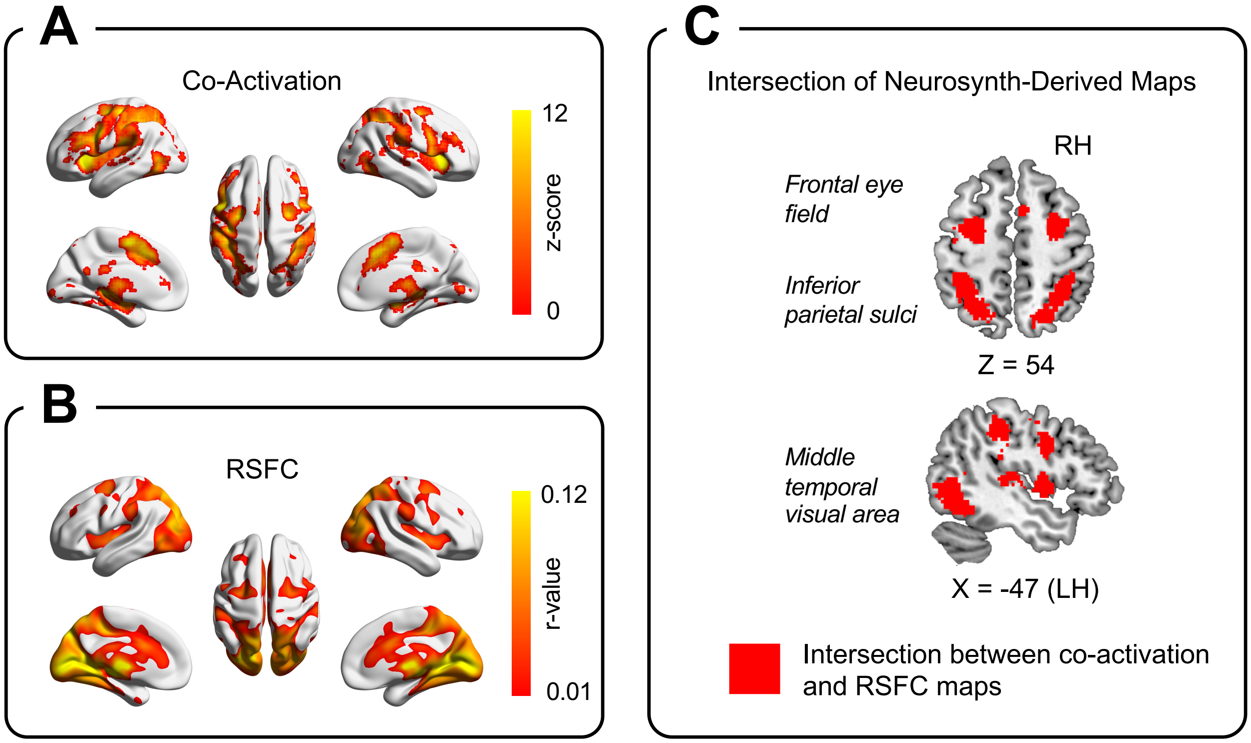


**Figure S3.** Brain maps of co-activation and RSFC produced by Neurosynth and their intersection. **A.** Brain map presenting regions co-activated with the PatAGE-cluster across more than 10,900 functional studies. A threshold of FDR corrected $q$-voxel < 0.01 was applied. **B.** RSFC from the PatAGE-cluster with the rest of the brain in the 1000 Functional Connectome dataset. A liberal cutoff value of $r$ = 0.01 as in the previous literature was used. **C.** Overlapping areas between co-activation and RSFC maps of the PatAGE-cluster. *Acronyms: FDR, false discovery rate; LH, left hemisphere; PatAGE-cluster, the cluster significantly associated with paternal age; RH, right hemisphere; RSFC, resting-state functional connectivity.*

**Table S1.** Zero-order correlations between familial variables, reading composite scores, and reading-related skills.

| Variable | 1 | 2 | 3 | 4 | 5 | 6 | 7 | 8 | 9 | 10 | 11 | 12 | 13 | 14 | 15 |
|---|---|---|---|---|---|---|---|---|---|---|---|---|---|---|---|
| 1 *t*1PA | — | | | | | | | | | | | | | | |
| 2 *t*1RAN | -0.125 | — | | | | | | | | | | | | | |
| 3 *t*2READ | **0.458** | **0.313** | — | | | | | | | | | | | | |
| 4 *t*2PA | **0.448** | -0.129 | -0.045 | — | | | | | | | | | | | |
| 5 *t*2RAN | -0.114 | **0.658** | 0.006 | -0.024 | — | | | | | | | | | | |
| 6 # Older Siblings | -0.288 | -0.174 | -0.248 | -0.155 | -0.243 | — | | | | | | | | | |
| 7 # Younger Siblings | 0.225 | 0.048 | **0.322** | 0.185 | -0.059 | **-0.596** | — | | | | | | | | |
| 8 PatAGE | -0.006 | -0.001 | **-0.385** | 0.017 | 0.263 | -0.113 | -0.223 | — | | | | | | | |
| 9 MatAGE | -0.045 | -0.049 | **-0.330** | -0.116 | 0.149 | 0.092 | **-0.346** | **0.633** | — | | | | | | |
| 10 PatARHQ | -0.153 | -0.202 | -0.294 | -0.008 | -0.003 | 0.194 | **-0.303** | -0.011 | 0.041 | — | | | | | |
| 11 MatARHQ | **-0.340** | 0.022 | **-0.464** | -0.056 | 0.042 | 0.105 | -0.118 | **0.336** | 0.197 | 0.051 | — | | | | |
| 12 PatEDU | 0.186 | -0.011 | 0.009 | 0.133 | 0.110 | **-0.310** | -0.052 | 0.226 | 0.168 | -0.227 | 0.104 | — | | | |
| 13 Mat EDU | 0.089 | 0.173 | -0.013 | -0.124 | 0.228 | **-0.353** | 0.065 | 0.192 | **0.379** | -0.047 | 0.155 | **0.442** | — | | |
| 14 SES | -0.002 | -0.006 | -0.260 | 0.002 | 0.261 | **-0.329** | -0.052 | 0.189 | **0.389** | -0.123 | 0.124 | **0.515** | **0.496** | — | |
| 15 HOME | 0.199 | 0.127 | **0.312** | -0.027 | 0.131 | -0.302 | **0.351** | 0.117 | -0.222 | -0.185 | -0.032 | -0.022 | -0.084 | -0.266 | — |

*Note:* ***Bold*** *text indicates a statistically significant correlation with a p-value less than 0.05.*
